# Dogs density drives the reproductive effort of American Oystercatchers (*Haematopus palliatus*) in disturbed habitats of the Maule Region, central Chile

**DOI:** 10.1101/2024.06.14.599056

**Authors:** Jesús Díaz, Fernando Medrano, Daniel Imbernón, Franco Villalobos, Juan Silva, Erik M. Sandvig, Sharon Montecino

## Abstract

The American Oystercatcher (*Haematopus palliatus*) is a shorebird specialized in coastal environments across the Americas. Thus, anthropic extensive use of the shoreline including the introduction of dogs and vehicles to beaches could directly impact fitness, and ultimately the persistence of the species’ populations. In this study, we aimed to assess the impact of pedestrians, vehicles and dogs on the nest density, productivity and hatching success of the American Oystercatcher in the Maule Region, central Chile. To this end, we sampled sandy beaches, quantifying the number of nests, eggs, chicks, but also pedestrians, vehicles and dogs during the breeding seasons of 2023 and 2024. We assessed for the influence of these threats on parameters of reproductive output by using Generalized Linear Mixed Models. We found that the number of dogs is the only variable that negatively impacts the number of nests, eggs and chicks of the American Oystercatcher in the sampled areas. Reducing the impact of dogs is a ubiquitous challenge in coastal environments of central Chile, however current regulations do not allow management relying on removal of dogs from important areas for biodiversity.

## Introduction

The South American Pacific coastline has been historically modified by humans (Rivera et al. 2021; Rivadeneira and Nielsen 2022). Currently, human impacts include the extraction of resources, habitat loss, and extensive use by recreational and commercial activities (Blanco-Libreros and Ramírez-Ruiz 2021; Day et al. 2021). Recreational uses include sun-and-beach tourism, dog walking, horseback riding, camping, fishing, and the transit of motorized vehicles. These activities usually restrict the habitats for biodiversity to sites with less human intervention (Senner et al. 2017).

Shorebirds are a group of species that are highly dependent on coastal environments. In the case of shorebirds, human-related disturbances can affect individuals’ breeding, roosting, and foraging activities (Palacios et al. 2022). Pedestrians, dogs, and motorized vehicle traffic on beaches also affect the energetic expenditure of shorebirds (Navedo et al. 2019; Cortés et al. 2021; Oliveros et al. 2021; Gómez-Serrano 2021). However, the ultimate effects on the demography of populations is a key response that is less understood, but is sorely needed for designing and implementing effective conservation actions.

The American Oystercatcher (*Haematopus palliatus*) is a widely distributed species of shorebird throughout the Americas. In the Atlantic coast, it ranges between the northern Atlantic coasts of the United States to southern Argentina, whereas on the Pacific coast it ranges from the northern tip of the Mexican coasts to the Chiloé archipelago in southern Chile (American Oystercatcher Working Group et al. 2020). The species is strictly constrained to coastal environments (O’Brien et al. 2006; Barros 2018), using sandy beaches and muddy estuaries, shell beaches, dunes, mudflats (O’Brien et al. 2006), intertidal zones, and salt production pools (American Oystercatcher Working Group et al. 2020). The American Oystercatcher is globally classified as Least Concern by BirdLife International (2016). However, in Chile it is locally classified as Near Threatened, due to loss of habitat and reported low reproductive success (RCE, 2020). The local population trends are poorly known, but breeding areas extensively overlap with unregulated tourism. Further, recent efforts to monitor breeding populations have reported that the species currently is experiencing low breeding success rates throughout Chile, yet the specific drivers have not been quantified (Montecino et al. unpublished data). This could be understood due to the fact that the species’ life history is characterized by delayed breeding, low reproductive rates, and long lifespans (Clay et al. 2014; American Oystercatcher Working Group et al. 2020), since they generally do not reproduce until they are 2-3 years of age (American Oystercatcher Working Group et al. 2020). It has also been described that ‘overwash’ from severe flooding/storm tides and mammalian predation are the leading causes of clutch failure before hatching (American Oystercatcher Working Group et al. 2020).

One of the areas with the highest reported densities of American Oystercatcher on the Pacific Coast of South America is the Mataquito River Mouth (Ortíz-Soazo et al. 2009), in central Chile. However, recent monitoring efforts report that the number of individuals in this area are declining (J. Díaz, unpublished data). In this study, we aim to understand how common anthropogenic threats affect the density of nests, productivity, and breeding success of the American Oystercatcher at this important river estuary.

## Methods

### Study area

We surveyed three areas of sandy beaches in the Maule Region: (i) Northern limit of the Mataquito River Mouth; (ii) La Trinchera beach, located between the Mataquito and Huenchullamí river mouths; and (iii) Beach next to the Huenchullamí River Mouth (Figure 1). In Mataquito and Huenchullamí river mouths, we conducted a ∼1km transect survey, and a 4.7km transect along La Trinchera beach. The dominant plant species are *Carpobrotus chilensis, Ambrosia chamissonis* and *Distichlis spicata*.

**Fig. 1.**
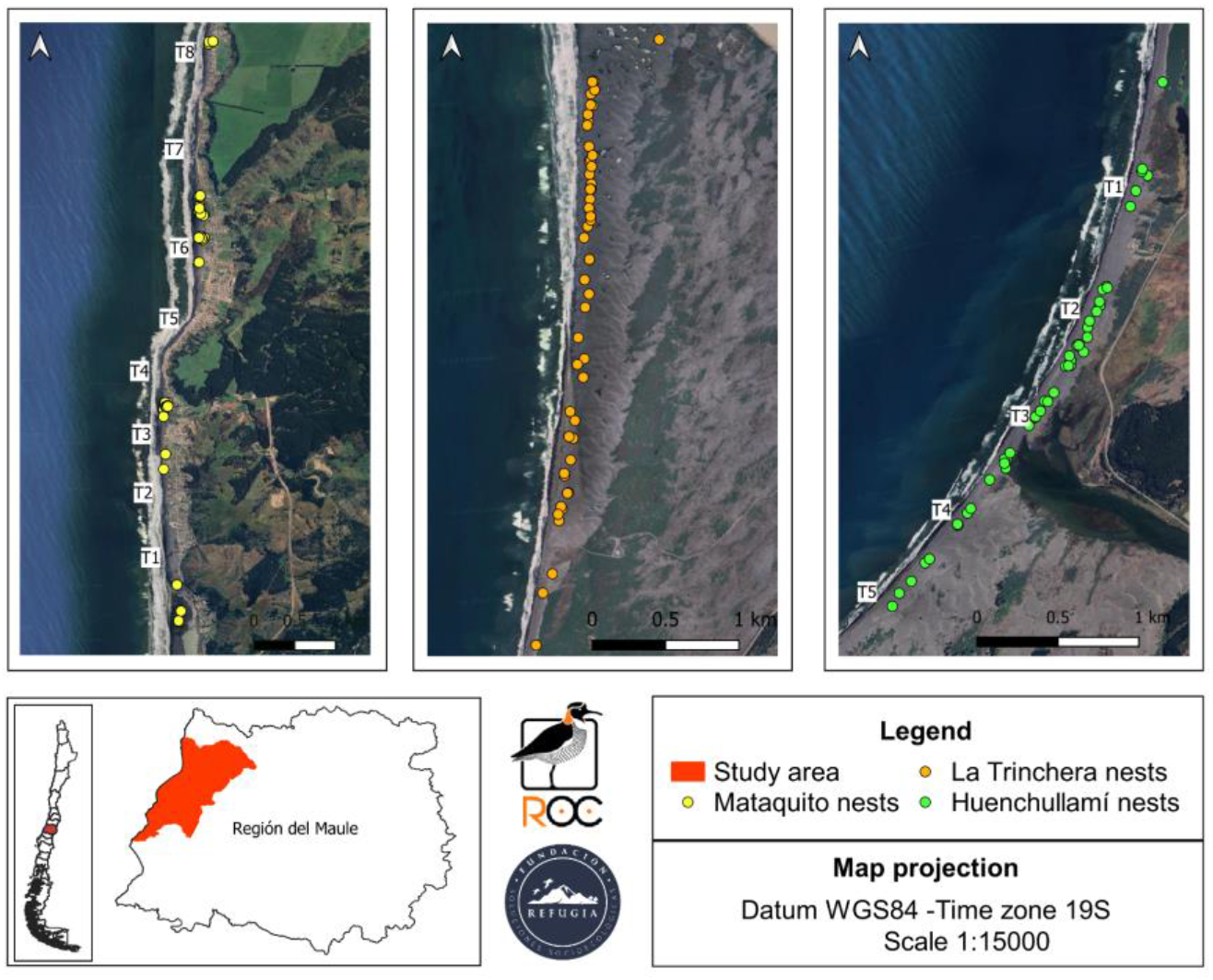
Map of the study area including (i), (ii) and (iii)

### Fieldwork

We surveyed each transect by foot, weekly, in two consecutive seasons. The first season spanned from January to May 2023, and the second season spanned from November 2023 to April 2024. In each site, we quantified the number of nests and within each nest, the number of eggs and chicks. We logged each nest using a GPS. During each visit and each transect, we quantified the potential threats in the areas, including the number of people encountered (pedestrians, fishermen, and beach-and-sun tourists), vehicles (cars, tractors, and motorbikes), and dogs (free-ranging or with owners but off-the-leash), as done by Palacios et al. (2022).

### Statistical analysis

We assessed the effect of the different threats on the number of nests through Generalized Linear Mixed Models (GLMM) using the *lme4* package in R (Bates et al. 2015). First, we built a model with the nest density (number of nests per km^2^) as the explanatory variable, the number of dogs, the number of pedestrians, and number of vehicles as fixed factors, and the year as a random factor. We built another model including the number of eggs + chicks (a proxy of productivity) as the explanatory variable, the number of dogs, the number of pedestrians, and number of vehicles as fixed factors, and the year as a random factor. In addition, we calculated the hatching success by calculating the ratio of chicks/eggs laid per nest. We built a model in which hatching success was the explanatory variable and the accumulated number of dogs, number of pedestrians, and number of vehicles in the whole season as fixed factors, and the year as a random factor. In both models, we assigned Gaussian or Gamma distributions depending on the residuals’ distribution of each variable. We did a backward selection, dropping variables and selecting the best models based on corrected AIC values.

## Results

### Nest density

The number of dogs negatively influenced the American Oystercatcher’s nest density (GLMM estimate: -0.09, df: 252, p-value: 0.002; Figure 2). The number of vehicles had no influence on the density of nests (GLMM estimate: 0.04, df: 252, p-value: 0582), nor did the number of pedestrians (GLMM estimate: 0.003, df: 252, p-value: 0.3381).

**Fig. 2.**
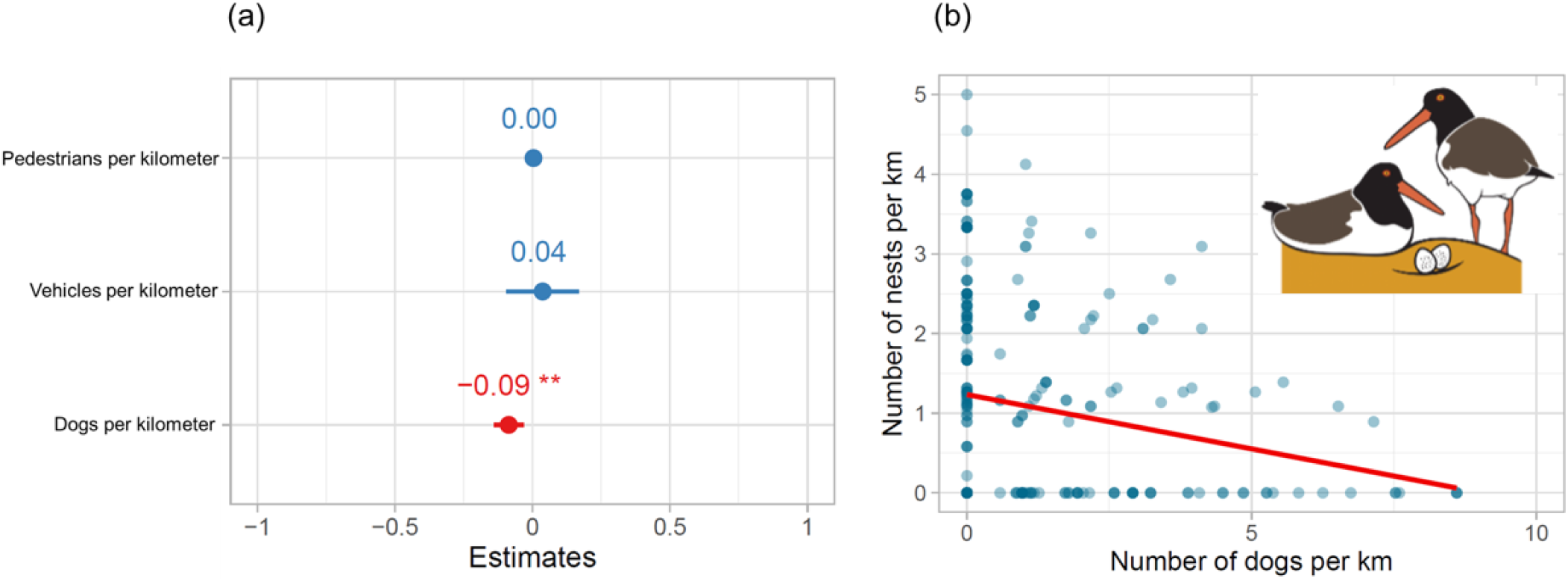
Drivers of the number of nests per kilometer of the American Oystercatcher in the Maule Region, coastal central Chile, including (a) Estimates of each anthropogenic driver on the number of nests per kilometer, and (b) Relationship between the number of dogs per kilometer and the number of nests of American Oystercatcher per kilometer. The red line shows the slope of the best fit of the linear model.Artwork by Felipe Cáceres

### Productivity

The number of dogs negatively influenced the American Oystercatcher’s number of eggs and chicks (GLMM estimate: -0.20, df: 252, p-value: 0.0017; Figure 3). The number of vehicles had no influence on the number of eggs and chicks (GLMM estimate: 0.07, df: 252, p-value: 0.6375), nor did the number of pedestrians (GLMM estimate: 0.005, df: 252, p-value: 0.4773).

**Fig. 3.**
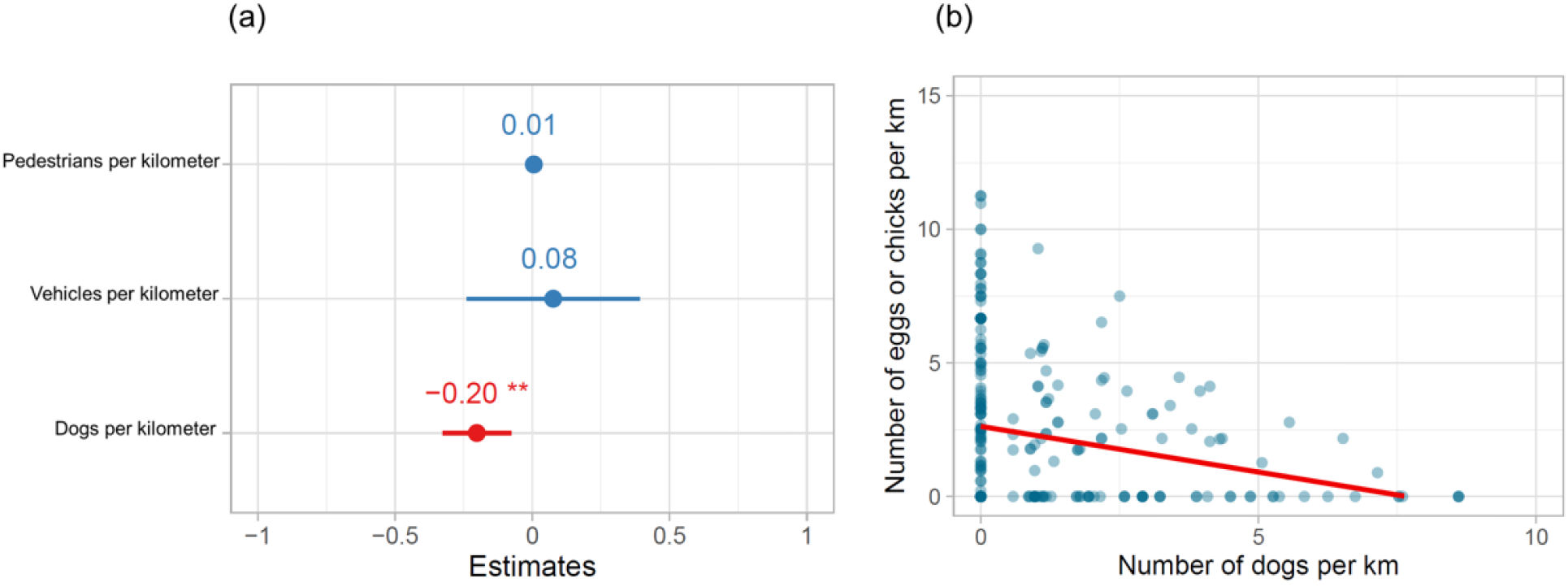
Drivers of the number of nests per kilometer of the American Oystercatcher in the Maule Region, coastal central Chile, including (a) Estimates of each anthropogenic driver on the number of nests per kilometer, and (b) Relationship between the number of dogs per kilometer and the number of eggs or chicks of American Oystercatcher per kilometer. The red line shows the slope of the best fit of the linear model.

### Hatching success

The accumulated number of dogs on the beaches did not negatively influence the hatching success of the American Oystercatcher (LME estimate: 0.0001, t-value: 0.052, p-value: 0.947), nor did the accumulated number of vehicles (LME estimate: 0.009, t-value: 0.845, p-value: 0.400), or the number of pedestrians (LME estimate: -0.0002, t-value: -0.513, p-value: 0.608) (Figure 4).

**Fig. 4.**
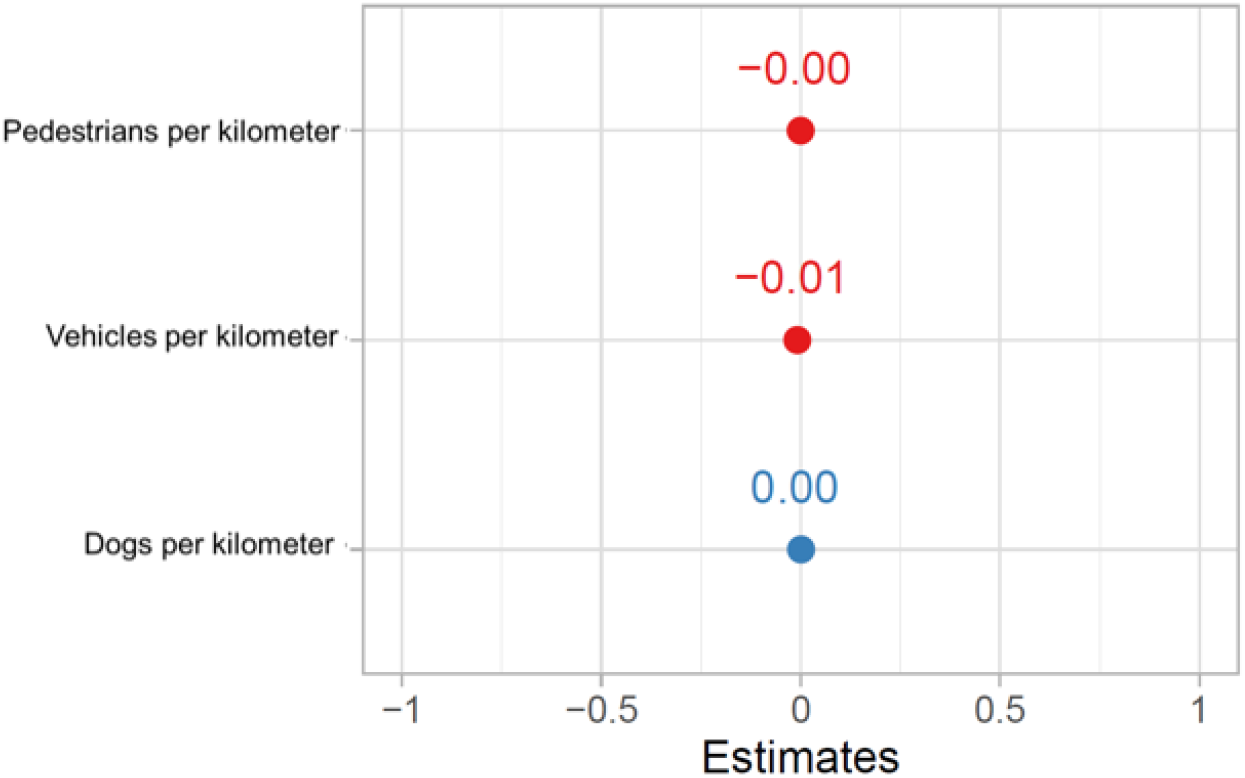
Drivers of the hatching success of the American Oystercatcher in the Maule Region, coastal central Chile, including estimates of each anthropogenic driver on the number of nests per kilometer

## Discussion

We found that the main driver of nest abundance and productivity is the number of dogs per kilometer on the beach, but that none of the anthropogenic drivers studied affect the hatching success. This might be explained by the American Oystercatchers avoiding areas with high levels of disturbances, which has been defined as “landscapes of fear” (Gaynor et al. 2019). Other shorebirds also avoid landscapes with high levels of disturbances (Gibson et al. 2021).

Dog management is complex and depends on the characteristics of the dog population. In Chile, free-roaming dogs are usually owned dogs with outdoor access (Contreras-Abarca et al. 2022; Silva-Rodríguez et al. 2023), which is likely to also be the case in our study area. Considering that the removal of dogs is forbidden in Chile, alternative management strategies are needed. In that context, some potential practices to avoid the impacts of dogs include (i) controlling the population size by keeping owned dogs inside properties (Silva-Rodríguez-et al. 2023); (ii) keeping dogs on-leashes on beaches, which has shown to reduce the negative effect on birds (Mengak et al. 2019); and (iii) creating ‘exclusion zones’ where the access to dogs is effectively controlled (Ferreira-Rodríguez et al. 2018; Pujol et al. 2022).

Although we did not find effects of the vehicles on the nest density and productivity, other studies in North America indicate that off-road vehicles directly impacts nest attendance of the American Oystercatcher (Felton et al. 2018), as well as daily nest survival and hatching success (Borneman et al. 2016). Furthermore, off-road vehicles have been shown to negatively affect the total abundance of some shorebirds (Tarr et al. 2010). Further information is needed to understand whether vehicles can affect other life history traits of shorebirds in central Chile.

Our manuscript shows that dog density directly reduces the number of nests and the productivity of an area. In addition, literature has widely shown that dogs increase the disturbances on shorebirds (Lilleyman et al. 2016; Navedo et al. 2019), and reduce the time dedicated to incubation (Gómez-Serrano 2021). Further research should be carried out to understand the long-term impact on the populations of the American Oystercatcher, and other species of shorebirds in coastal South America. For doing so, it is important to conduct and standardize banding efforts to understand other demographic parameters such as juveniles and adult survival in coastal landscapes that are currently dominated by dogs.

## Acknowledgments

Fieldwork was funded by the Coastal Solutions Fellowship Program of the Cornell Lab of Ornithology, David and Lucile Packard Foundation, Red de Observadores de Aves y Vida Silvestre de Chile (ROC), The Nature Conservancy, Manomet Inc., and Neotropical Migratory Bird Conservation Act.

